# The FERM Guild: A Differentially Correlated Microbial Module Drives Hypertension via Metabolic Flux Perturbations

**DOI:** 10.64898/2025.12.08.693055

**Authors:** Wenkai Lai, Yuchen Zhang, Shaoping Huang, Shirong Lai, Fuxin Lin, Ziwei Wang, Shanwen Sun, Fenglong Yang

**Author notes:** These authors contributed to the work equally and should be regarded as co-first authors (Wenkai Lai, Yuchen Zhang).

## Abstract

**Background:** Hypertension is a major risk factor for cardiovascular diseases, with changes in gut microbiota composition and function being closely associated with its onset and progression. However, the high inter-individual variability in gut microbiota complicates the identification of pathogenic mechanisms using traditional methods. In contrast, the smaller variability in gut microbial metabolites offers a more reliable and consistent basis for cross-individual comparisons.

**Results:** Parsimonious Flux Balance Analysis (pFBA), integrated with Double Machine Learning (DoubleML), identified 17 metabolites significantly associated with hypertension (*p* < 0.05, *RobustnessV alue* > 0.1). These included meso-2,6-diaminoheptanedioate, p-hydroxyphenylacetic acid, cellobiose, dextran 40 (1,6-*α*-D-glucan), L-glutamic acid, kestopentaose, among others. Differential microbial correlation network analysis identified a key microbial sub-network, the FERM guild, consisting of 19 species, with prominent genera including *Faecalibacterium,Enterobacter,Roseburia*, and *Methanobrevibacter*. Using GSEA, the dysregulation of this guild was found to be strongly associated with a set of 17 hypertension-related metabolites(*p* = 0.017). Further analysis revealed that the contribution of FERM genus to key metabolites is significantly associated with blood pressure (*p* < 0.05), even without significant differences in their abundance; additionally, an imbalance exists between FERM genus and other species.

**Conclusions:** Our findings reveal that hypertension is associated with a disruption of gut microbial diversity, structure, and metabolic function. Seventeen key metabolites related to blood pressure regulation were identified, exhibiting pro- or anti-hypertensive potential and linked to functional microbial modules. These results highlight the gut microbiota and its metabolites as promising targets for therapeutic intervention in hypertension.

## Introduction

Hypertension remains one of the major risk factors for cardiovascular diseases (CVD) such as stroke and heart failure. In addition, it is an important associated factor for common comorbidities such as chronic kidney disease, obesity, and type 2 diabetes[36, 22]. In 2010, approximately 31% of the global population had hypertension, making it a global public health issue [25]. Despite extensive research and interventions, blood pressure control still faces many challenges [45].

Recent studies have shown that the gut microbiota plays a crucial role in the development of cardiovascular diseases [43]. For instance, patients with heart failure with preserved ejection fraction (HFpEF) exhibit significant gut dysbiosis, with marked differences compared to healthy individuals [4]. The gut microbiota metabolizes dietary choline, phosphatidylcholine, and L-carnitine to produce trimethylamine (TMA), which is further oxidized to trimethylamine N-oxide (TMAO), a metabolite known to promote atherosclerosis [20, 43]. These findings provide new perspectives and insights into the etiology of hypertension and potential therapeutic strategies.

The gastrointestinal tract is the largest immune cell compartment in the body, representing the intersection between the environment and the host [27]. Lifestyle can shape and be influenced by the microbiome [50, 35, 46], thereby altering the risk of developing hypertension. A well-studied example is the consumption of dietary fiber, which leads to the production of short-chain fatty acids and contributes to the expansion of antiinflammatory immune cells, thus preventing the progression of hypertension [2]. The gastrointestinal tract is also the primary site for the interactions between dietary components, gut microbiota and their metabolites, and antihypertensive drugs, with blood pressure potentially regulated by gastrointestinal hormones [42].

Although species-level analysis can provide basic information about microbial composition, it does not fully capture the complexity and functional diversity of microbial communities. Species-level analysis overlooks interactions between microorganisms and their functional potential. Currently, research on hypertension-related gut microbiota is relatively limited. Traditional microbial studies often focus on changes in microbial species abundance, such as *α*-diversity, *β*-diversity, and species abundance differences [47]. However, challenges in microbial research include data sparsity and significant inter-individual variability, which complicate and reduce the accuracy of constructing risk assessment or sample classification models. The interaction networks between microorganisms are crucial for maintaining gut health and metabolic balance [37, 13], yet these interactions may not be fully captured at the species level. The impact of gut microbiota on the host is typically mediated through interactions between their metabolites and the host. [18, 1]. Compared to simple species analysis, metabolomics has many advantages in deciphering the composition and function of microbes within an organism: Gut microbiota metabolites, as signaling molecules and substrates for host metabolic responses, influence various physiological and pathological processes in the host [19]. For example, short-chain fatty acids (SCFAs) are one of the most studied classes of small molecule metabolites. They are produced by gut microbes through the fermentation of dietary fibers and can regulate host physiological and biochemical functions. These functions include maintaining the natural intestinal barrier at the colonic epithelium and mucus levels, regulating gut motility, secreting gut hormones, modulating chromatin, influencing the gut-brain axis, and supporting immune functions. Metabolites are the end products of metabolic activities in the body, and their composition and abundance directly reflect the physiological state, metabolic pathways, and levels of biological activity within the organism [34, 15]. Metabolomics also offers strong biological interpretability. Metabolites are often closely associated with physiological processes and metabolic pathways in the body [44], which endows metabolomic data with significant biological relevance and interpretability.

Recent studies have shown that hypertension is closely related to disturbances in the metabolites of the gut microbiota [49]. The gut microbiota can influence host health, particularly through its metabolites. These metabolites can enter the bloodstream and affect the physiological and pathological states of the host. For example, some studies have found that certain gut microbial metabolites, such as SCFAs, can counteract hypertension by modulating immune responses, reducing inflammation levels, regulating energy balance and cholesterol metabolism [41]. Additionally, some gut microbiota-derived metabolites may be involved in blood pressure regulation, such as TMAO produced from dietary nutrients by gut microbes, which has been associated with poor cardiovascular health, including atherosclerosis, hypertension, and heart disease [20]. In this study, parsimonious Flux Balance Analysis (pFBA) was used to calculate the metabolic flux spectrum for each sample. Metabolic flux reflects the rate at which microorganisms produce and consume metabolites, which is related to the state of the host’s disease. This research explore the relationship between metabolites and disease phenotypes, ultimately identifying specific species. We systematically organized microbiome data for comprehensive analysis, constructing a knowledge-based microbial cross-feeding network. This enabled a systematic analysis of the interactions between microbes and metabolites and their relationship with the occurrence of hypertension.

## Materials and methods

### Overview of the Study Workflow

In this study, 16S rRNA gut microbiome sequencing data related to hypertension, along with corresponding clinical phenotype information, were downloaded from the European Nucleotide Archive (ENA) database. Samples were classified into hypertension and healthy control groups according to the World Health Organization (WHO) blood pressure diagnostic criteria (Figure 1A). Subsequently, *α*-diversity and *β*-diversity analyses were performed, and microbial taxa with statistically significant differences were identified (Figure 1B). Using microbial species information recorded in the AGORA2 metabolic model database, samples with low matching fidelity to the database were excluded. The retained samples were then used to infer metabolic fluxes based on parsimonious Flux Balance Analysis (pFBA), and a cross-feeding network at the microbial community level was constructed accordingly (Figure 1C). Integrating clinical blood pressure measurements with medium metabolic flux features (net influx or efflux in the community model), a Double Machine Learning (DoubleML) framework was applied to control for potential confounders. This enabled estimation of the average treatment effects of individual metabolites on systolic and diastolic blood pressure, followed by sensitivity analyses to identify robust candidate metabolites (Figure 1D). Building on this, Gene Set Enrichment Analysis (GSEA) was employed to evaluate the enrichment of key metabolites within distinct microbial clusters (Figure 1E). To further investigate the dysbiosis of key guilds, this study extracted the key metabolites and species interactions from the cross-feeding network. We performed association analysis between these interactions and blood pressure in both the discovery and validation cohorts. Additionally, we conducted ADBA distal balance analysis on the species within key guilds to validate their dysbiosis (Figure 1F). Finally, the production and consumption rates of metabolites at the community level were quantified and their associations with blood pressure phenotypes were analyzed after adjustment for confounders using DoubleML. Metabolic pathway enrichment analysis was further conducted on metabolites showing significant associations to elucidate their potential physiological functions and mechanisms (Figure 1G).

**Fig. 1.**
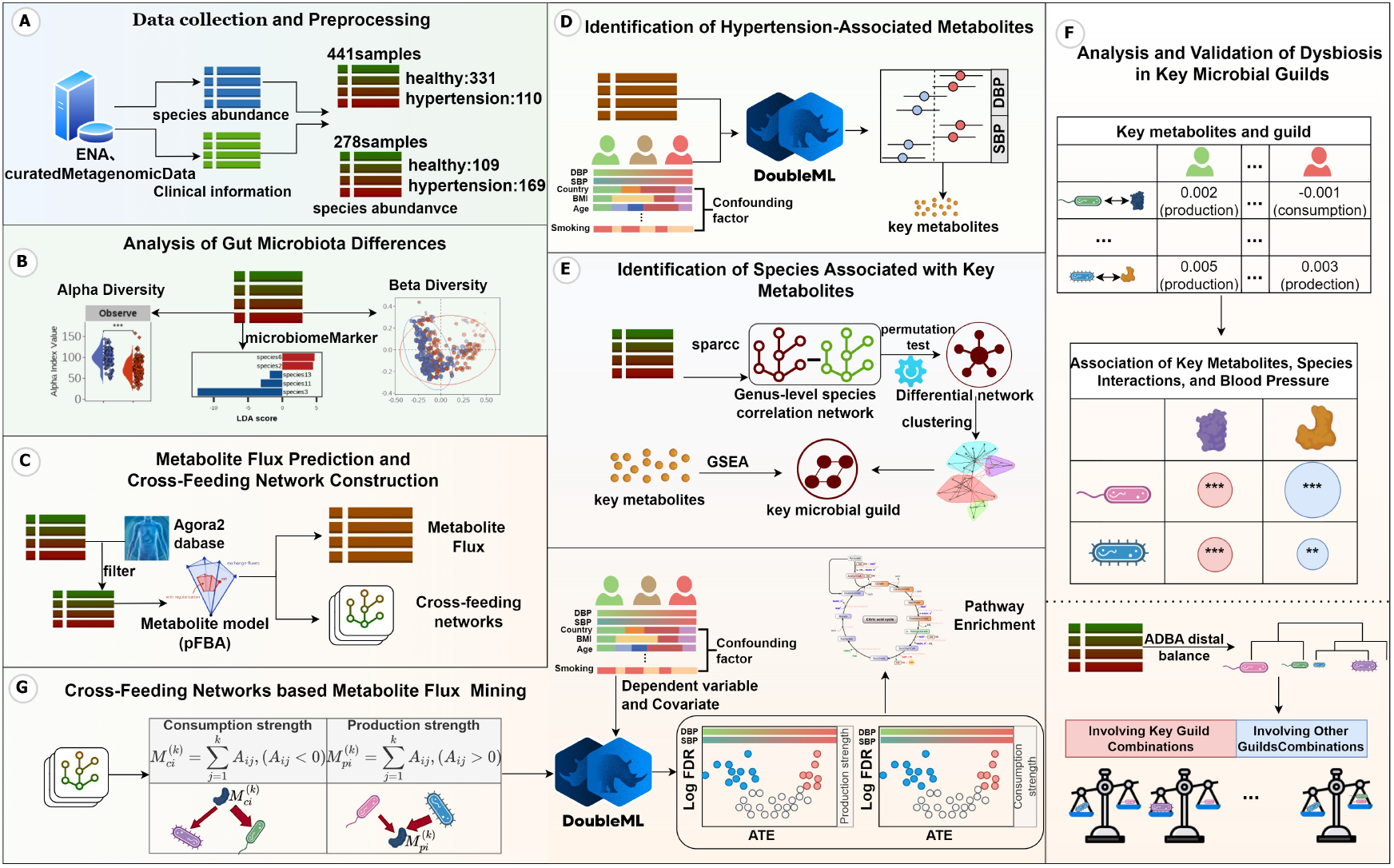
Overview of the research workflow. Downloading and preprocessing of hypertension gut microbiota data (Step A), analysis of gut microbial differences (Step B), prediction of metabolic flux based on microbial data (Step C), identification of Hypertension-Associated Metabolites (Step D), identification of species associated with key metabolites (Step E), Analysis and validation of dysbiosis in key metabolites guild (Step F). Cross-Feeding Networks based Metabolite Flux Mining (Step G)

### Data Collection and Sample Grouping

In this study, species abundance tables derived from 441 fecal microbiota samples (16S rRNA sequencing) were obtained from the work of Jacobo de la Cuesta-Zuluaga et al [14], available at https://github.com/jsescobar/westernization/tree/master/files. Each sample was accompanied by detailed clinical information, including systolic and diastolic blood pressure, age, gender, body mass index (BMI), cholesterol, low-density lipoprotein (LDL), cardiometabolic disease status, and medication use. According to the World Health Organization (WHO) definition of hypertension (diastolic blood pressure ≥ 90 mmHg or systolic blood pressure ≥ 140 mmHg), participants were classified into a hypertension group (*n* = 110) and a healthy control group (*n* = 331) for downstream analyses. Additionally, we collected 169 hypertension samples from curatedMetagenomicData, a specialized database for human gut microbiome data. Since the age distribution of the hypertension samples ranged approximately from 55 to 70 years, and the samples were primarily from Austria (AUT), China (CHN), and Italy (ITA), we established matching criteria for healthy controls to minimize confounding effects. The selection criteria for healthy samples were as follows: age between 55 and 70 years, country of origin limited to AUT, CHN, or ITA, and a BMI between 18.5 and 23.9. Based on these criteria, 109 healthy samples were obtained, resulting in a total of 278 samples. The species abundance data for these samples can be retrieved in R using the corresponding sample names provided in Table S1. This dataset was used for validating the statistical analysis results.

### Identification of Hypertension-Associated Metabolites

To investigate the relationship between metabolic fluxes and hypertension, we first filtered the metabolites by extracting the community-level metabolites (metabolites representing the overall import and export fluxes between the microbial community and the external medium). We then retained only those metabolites with nonzero predicted fluxes in at least 10% of the samples. Given that the underlying mechanisms leading to elevated diastolic and systolic blood pressure may differ, we conducted separate analyses to identify metabolites associated with diastolic blood pressure (DBP) and systolic blood pressure (SBP). To account for potential confounding factors, we included Age, Adiponectin levels, BMI, Body Fat Percentage, Fasting Glucose, Glycated Hemoglobin (HbA1c), High-Density Lipoprotein (HDL), High-Sensitivity C-Reactive Protein (hs-CRP), Insulin levels, Total Cholesterol, Triglycerides, Very Low-Density Lipoprotein (VLDL), Waist Circumference, Residential Location (city), Gender, Smoking Status and Medication Use as covariates in the models. Using the Double Machine Learning (DoubleML) framework with double random forests [11], we built separate models for DBP and SBP and estimated the average treatment effect (ATE) of each metabolite on these two measures of blood pressure. Each metabolic flux was treated as the treatment variable, with DBP)or SBP as the outcome variable, and the average treatment effect (ATE) on blood pressure was estimated within the DoubleML framework. Since both the treatment and outcome variables are continuous, DoubleML employs a residualization step using RandomForestRegressor to fit two necessary conditional expectation functions: *m*(*X*), representing the relationship between the metabolic flux and the covariates, and *g*(*X*), representing the relationship between the outcome variable and the covariates. Causal effects of the metabolic fluxes on blood pressure were then estimated after residualization, effectively removing the influence of confounding factors. To fit the ATE for each metabolite, the random forest regressors were used with default hyperparameters and a 5-fold cross-validation procedure. Furthermore, p-values for each metabolite were adjusted using the false discovery rate (FDR) correction. To assess the potential impact of unobserved confounders, we assumed a correlation of 0.1 between the unobserved confounders and both the exposure (*cf_d*) and the outcome (*cf _y*). Sensitivity analyses were performed to calculate the robustness values (RV) for each metabolite [12], and metabolites with RV > 0.1 were considered as hypertension-associated metabolites.

### Networks based Mining of Network topology Characteristics

#### Microbial Correlation Differential Network Analysis

Gut microbiota can influence host health, particularly through their metabolic products. Since gut microorganisms often function cooperatively, we used the microbial species abundance matrix to calculate the correlations between species using the SparCC method. We then constructed a microbial correlation network *G*_*hypertension*_ and *G*_*healthy*_ by sparsifying the matrix using the overall mean correlation as a threshold. To calculate the microbial correlation differential network *△G*, we subtracted the edge weights of the healthy microbial network from those of the hypertensive microbial network.

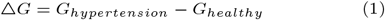

We then used the permutation test [16] to assess the significance of the edge differences between the healthy and hypertensive networks, filtering out non-significant edges.

#### Clustering and Enrichment of Microbial Nodes in the Differential Network Related to Metabolic Important Features

To explore which microorganisms mainly contribute to significant metabolic fluxes, this experiment calculated the differential correlation network of microorganisms and used a multi-level modularity optimization algorithm [6] to cluster the nodes. Subsequently, the Gene Set Enrichment Analysis (GSEA) method [38, 26] was used to evaluate the enrichment of each functional cluster in key metabolic features. Specifically, microbial species were ranked based on the stability of their associations with key metabolites. The stability metric *S*_*i*_ was calculated as follows:

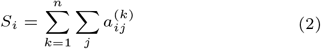

where *S*_*i*_ represents the overall stability of the association between microbial species *i* and the set of key metabolites; *k* denotes the *k* − *th* sample; *j* denotes the *j* − *th* key metabolite; and 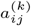 refers to the element of the binary adjacency matrix in the cross-feeding network in sample *k*.

Subsequently, microbial species were ranked in descending order based on their *S*_*i*_ values, and GSEA was applied to identify microbial guilds that are highly associated with the regulation of key metabolite homeostasis.

### Analysis and Validation of Dysbiosis in Key Microbial Guilds

To clarify the differential role of key guilds in regulating critical metabolic fluxes, we first extracted the associations between the key guild and key metabolites from the overall network. In the discovery cohort, the DoubleML method was applied to evaluate the independent contribution of each microbe–metabolite relationship to systolic and diastolic blood pressure, consistent with the approach used for key metabolite identification. Since the validation cohort phenotype was categorical without specific blood pressure measurements, only Wilcoxon tests were conducted on these relationships to identify microbe–metabolite associations related to hypertension.

Furthermore, to assess the balance state of key guilds in the gut microbiome, the ADBA distal balance method[31] was employed to identify balance combinations that best distinguish the two groups. Wilcoxon tests were then performed on these identified balance combinations to screen for those with significant differences.

## Results

### Hypertension-Associated Alterations in Gut Microbiota

In this study, gut microbial data were divided into two groups based on disease status: hypertensive and healthy. *α*-diversity analysis on the gut microbiota of both the hypertensive and healthy groups was conducted, six indices: Observe, Chao1, ACE, Shannon, Simpson, and Pielou were calculated. Observe, Chao1 and ACE showed significant differences, with the healthy group exhibiting higher values than the hypertensive group (Figure 2A), indicating a decrease in microbial richness in hypertensive patients while evenness showed no significant difference.

**Fig. 2.**
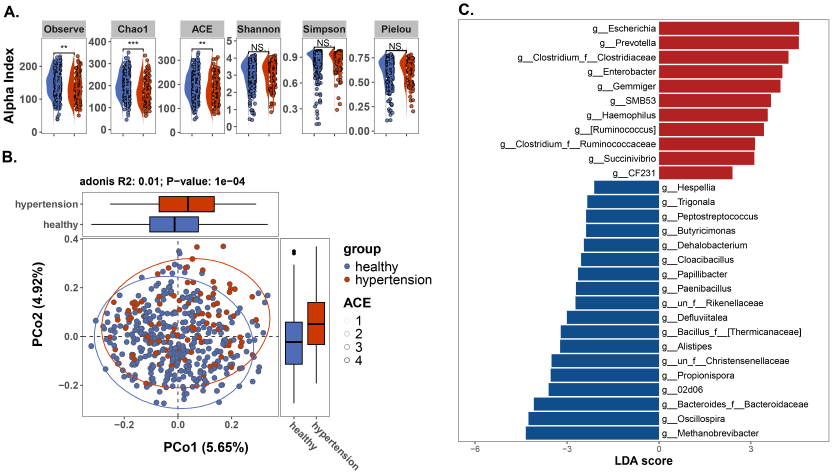
Hypertension-Associated Alterations in Gut Microbiota: Wilcoxon rank-sum test revealed significant differences in the Observe, Chao1 and ACE *α*-diversity indices between hypertensive and healthy individuals. Red indicates hypertensive patients, while blue denotes healthy controls (A). significant gut microbial composition differences between the hypertensive and healthy samples (B). A larger proportion of gut microbial species exhibit reduced abundance in individuals with hypertension (C).

The *β*-diversity analysis based on Bray-Curtis distance was conducted to assess differences in gut microbial community structure between the hypertension and healthy groups. The analysis showed that group status explained approximately 1% of the total variance (R^2^ = 0.01). The permutation test indicated that the difference was statistically significant (*P* = 10^−4^) (Figure 2B).

Using linear discriminant analysis (LDA), we identified a total of 29 genera that showed significant differences between the hypertension and healthy groups, with 11 genera enriched in the hypertension group and 18 genera enriched in the healthy group (Figure 2C). When extending the LDA across all taxonomic levels (kingdom, phylum, class, order, family, genus, species) without restriction, we detected 64 taxa enriched in the healthy group and 32 taxa significantly enriched in the hypertension group (Figure S1,Table S2). Overall, these findings indicate that the gut microbial abundance in hypertension patients tends to be reduced, consistent with the decreased species richness observed in the previous *α*-diversity analysis. However, from an overall perspective, the *β*-diversity analysis showed that the variance explained by the principal coordinates in the PCoA, as well as the *R*^2^ value from the PERMANOVA analysis, were both low, indicating limited differences in microbial community structure. Therefore, it is scientifically reasonable and necessary to further investigate the system from the perspective of metabolic flux and microbial interspecies interactions.

### Altered Community-Level Medium Metabolite Fluxes in the Gut Microbiota of Hypertensive Patients

Analyzing species abundance data alone cannot fully capture the complexity and functional variability of microbial communities, and it may overlook interactions and functional potential among microbes. Therefore, metabolic fluxes were predicted and systematically evaluated to reveal their potential associations with hypertension. To enable accurate modeling of gut microbiota metabolic fluxes, the detected microbial species were mapped to the AGORA2 database. Samples in which the cumulative relative abundance of the mapped species fell below 80% were excluded. Following this filtering step, 312 samples were retained, including 226 from healthy individuals and 86 from patients with hypertension(Figure 3A). Metabolic flux distributions were inferred using parsimonious Flux Balance Analysis (pFBA), allowing the systematic estimation of reaction-level metabolic activity within the microbial community.

**Fig. 3.**
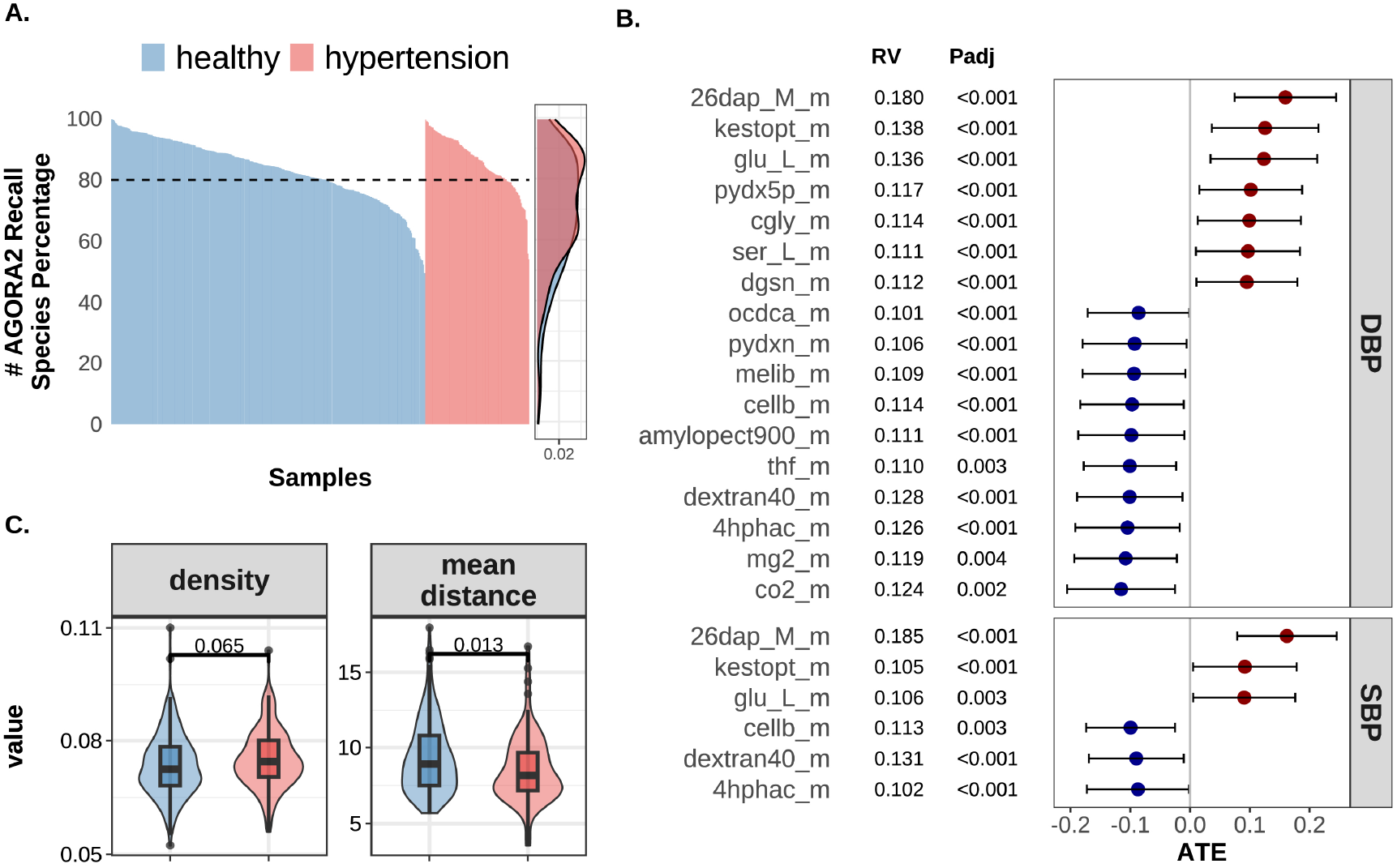
Altered Community-Level Medium Metabolite Fluxes in the Gut Microbiota of Hypertensive Patients: Samples were filtered based on a matching threshold of less than 80% with the AGORA2 database. The height of each bar represents the cumulative relative abundance of microbial species in the sample that successfully matched the AGORA2 reference models. Red bars indicate hypertensive patients, and blue bars represent healthy individuals (A). Seventeen metabolites exhibited significant average treatment effects on diastolic blood pressure, while six metabolites showed average treatment effects on systolic blood pressure. Red indicates a positive average treatment effect, and blue indicates a negative average treatment effect. RV represents the robustness value derived from sensitivity analysis (B). The average shortest path length of the cross-feeding network for 17 metabolites is significantly reduced in hypertension (C).

Using Double Machine Learning (DoubleML) analysis, we identified 17 metabolites significantly associated with diastolic blood pressure (DBP), among which 7 metabolites showed a positive average treatment effect (*ATE* > 0) and 10 showed a negative ATE (*ATE* < 0). Additionally, 6 metabolites were found to be significantly associated with systolic blood pressure (SBP), with 3 metabolites having *ATE* > 0 and 3 having *ATE* < 0. Notably, the six systolic blood pressure-associated metabolites completely overlapped with those identified for diastolic blood pressure. Statistical analysis showed that the false discovery rate (FDR) for these metabolites was below 0.005. Sensitivity analyzes, performed under the assumption of an unobserved confounder strength (*cf y* and *cf d*) set at 0.1, yielded robustness values exceeding 0.1, indicating the stability of these associations (Figure 3B,Table S3).

For each sample, cross-feeding networks were constructed, and subnetworks composed of hypertension-associated metabolites were extracted. Network density and average shortest path length were calculated. No significant difference was observed in network density between the hypertension and healthy groups (P = 0.065); however, the average shortest path length was significantly lower in the hypertension group compared to the healthy group (P = 0.013), These findings suggest that although the number of species contributing to the hypertension-associated metabolites did not significantly differ between groups, the rates at which these metabolites were produced or consumed by the microbial community were significantly elevated in hypertension (Figure 3C).

### Dysregulation of Community-Level Medium Metabolic Flux Driven by a Functional Group Disturbance

The multi-level modularity optimization algorithm was employed to cluster nodes in the differential microbial interaction network, and identifying a total of 9 guilds(Figure 4A). Based on the shared 17 key metabolic flux-associated species identified by DoubleML, species were ranked according to the stability of the metabolite-species associations. Using the GSEA method, each guild was analyzed for enrichment of important metabolic flux-associated species. It was found that nodes in the third guild ranked significantly higher in terms of frequency among the important metabolic flux-associated species, with most of the rankings within the top 50(Figure 4B,Table S4). In guild 3, there were severe disturbances in the interaction relationships at the genus level, particularly among species such as *Faecalibacterium, Enterobacter, Roseburia*, and *Methanobrevibacter* (FERM). These interactions showed significant differences. From the gutMDisorder database [30], it was found that species within this guild, *Roseburia* have already been reported to be associated with hypertension. The differential network composed of the FERM guild exhibited significant changes in edge weights (p < 0.05), indicating alterations in the strength of interspecies interactions (Figure 4C). Therefore, it is inferred that altered interaction patterns within FERM can lead to abnormalities in the 17 important metabolites and ultimately influence hypertension.

**Fig. 4.**
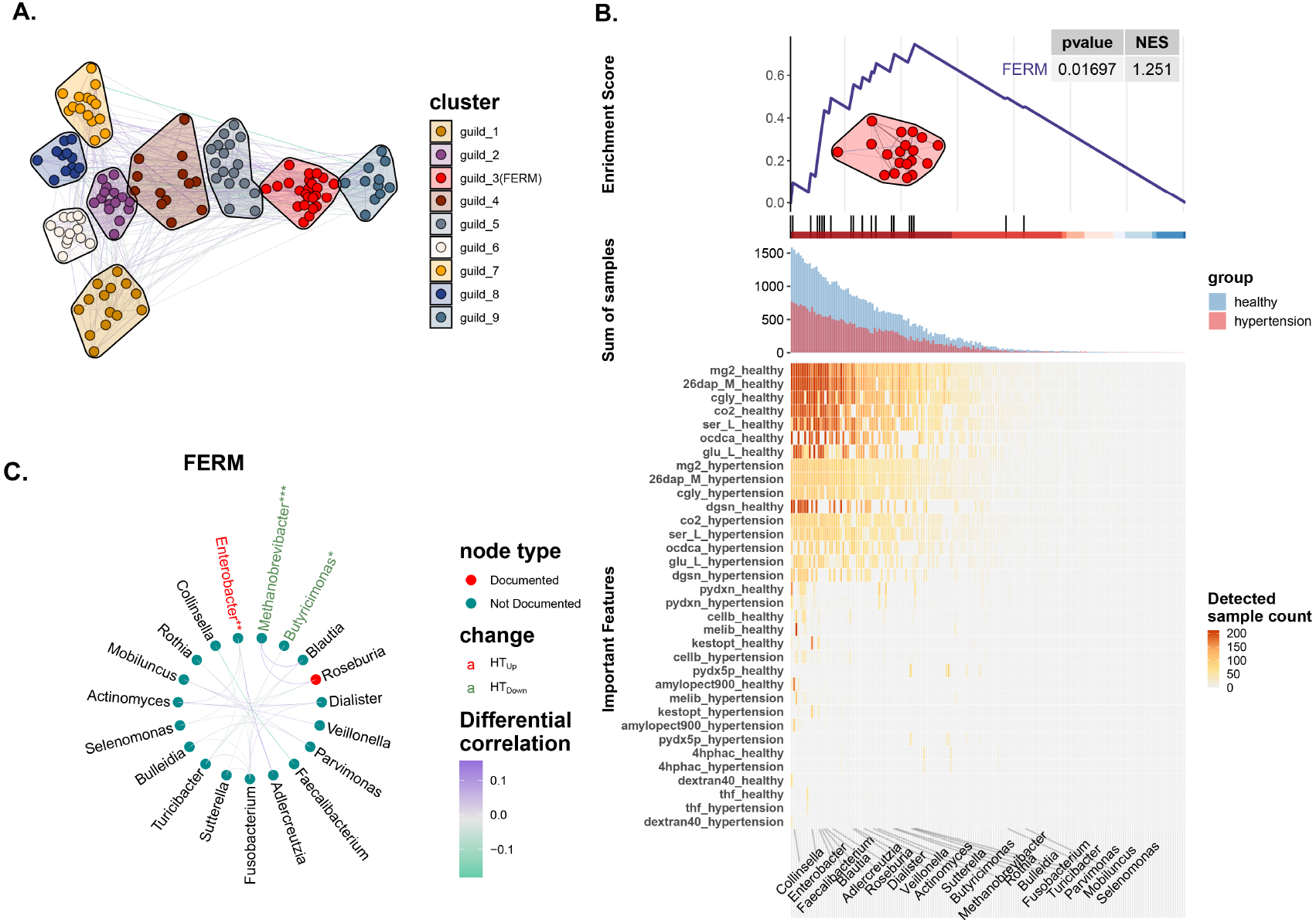
Dysregulation of Community-Level Medium Metabolic Flux Driven by a Functional Group Disturbance: The microbial correlation difference network was divided into 9 guilds using multi-level modularity optimization algorithm (A). The GSEA algorithm was used to analyze the enrichment of the 17 key metabolites in microbial guilds. The 17 key metabolites are primarily associated with FERM. NES is the normalized enrichment score. The color of the heatmap below represents the number of samples in which each species is associated with the 17 key metabolites (B). The microbial interactions in the FERM have undergone significant changes and a red species in the FERM have already been shown to be associated with hypertension, green edges indicates a strengthened negative correlationtions, and purple edges indicates a strengthened positive correlation (C). .

### Significant Association Between the Principal Eigenvector of the FERM Guild and Blood Pressure

Additionally, we extracted the principal eigenvector for each functional guild and fitted linear regression models with blood pressure as the outcome. The results showed that the principal eigenvector of FERM was significantly associated with DBP (coefficient = 30.4, p = 0.006) and SBP (coefficient = 40.9, p = 0.0129). In contrast, the remaining guilds showed few significant associations with blood pressure (Figure S2A,B; Table S5). In an independent validation cohort of 278 samples, the principal eigenvector of FERM also differed significantly between the hypertensive and healthy groups (Wilcoxon rank-sum test, p = 0.0012) (Figure S2 C.).

### FERM Guild Exhibits Differential Regulatory Contributions to Key Metabolic Fluxes

To assess differential contributions of the FERM guild to key metabolic fluxes, we extracted its associations with key metabolites and applied DoubleML to evaluate their independent effects on blood pressure. In the discovery cohort, 17 FERM species showed significant flux differences involving 13 metabolites, with six positive and seven negative associations with blood pressure (Figure 5A). In the validation cohort, 11 species–9 metabolite associations were significant, including five positive and four negative (Figure 5B). Across cohorts, 11 species–metabolite links were consistently replicated (Figure 5C), indicating robust regulatory roles of specific FERM members in blood pressure–related metabolic fluxes.

**Fig. 5.**
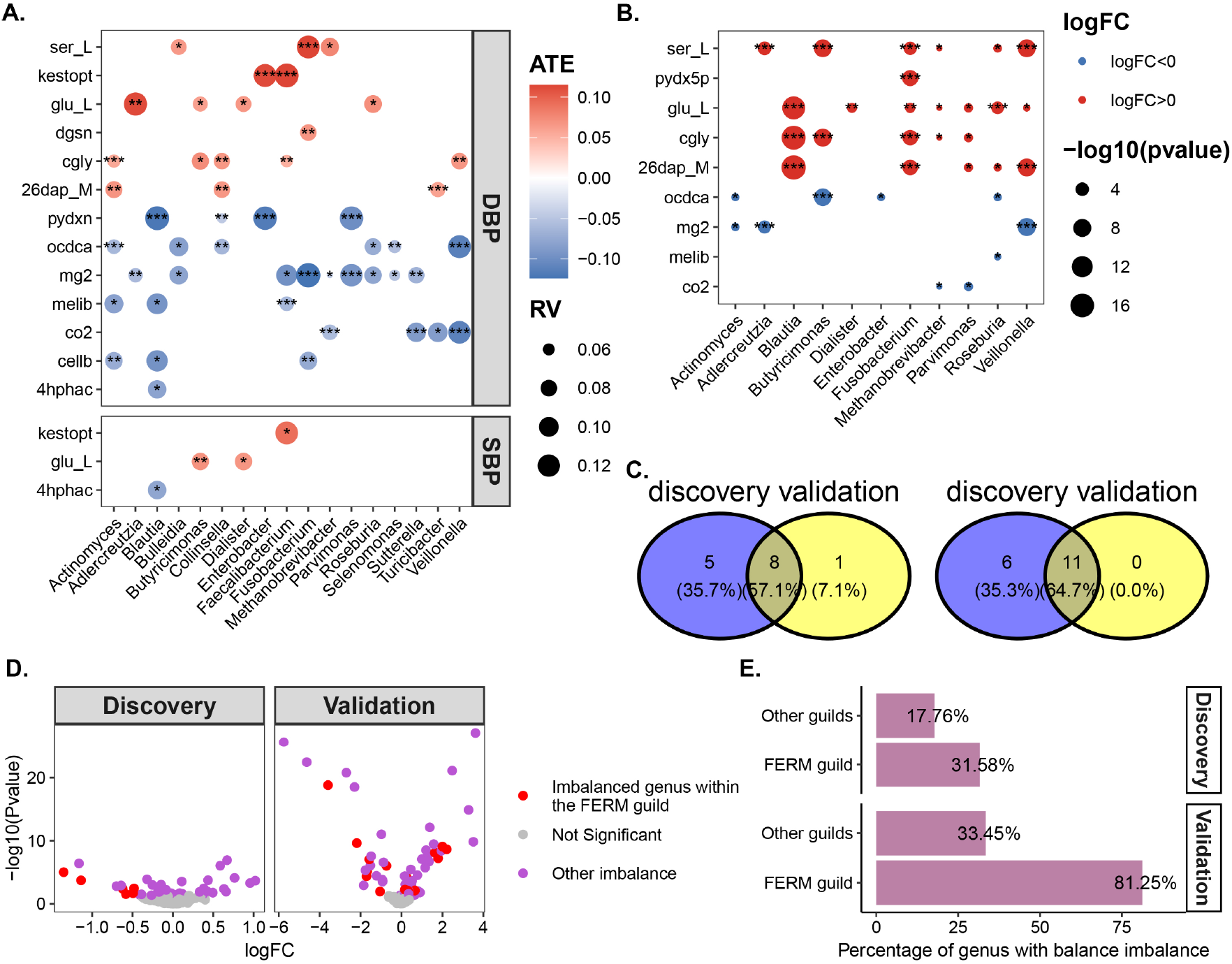
Metabolic Flux and Inter-species Relationship Imbalance in the FERM Guild: The contribution of FERM Genus to key metabolites and its association with blood pressure in the discovery cohort, with red bubbles indicating positive correlation and blue bubbles indicating negative correlation (A). The contribution of FERM Genus to key metabolites and its association with blood pressure in the validation cohort, with red bubbles indicating positive correlation and blue bubbles indicating negative correlation (B). Intersection of key metabolites and FERM Genus in the discovery and validation cohorts (C). Differential analysis of balance relationships identified by the ADBA distal balance method. Gray indicates no significant difference, red indicates significant differences involving the FERM genus, and blue indicates significant differences involving other genera (D). Statistical analysis of balance relationships involving the FERM genus and other genera (E).

### Pronounced Systemic Imbalance Among FERM Species

To comprehensively evaluate imbalance within the FERM guild, we computed species balances using the sequential binary partition (SBP) framework. In the discovery cohort, 40 significantly imbalanced balances were identified, six of which involved FERM species. In the validation cohort, 63 significant balances were detected, including 13 associated with FERM species (Figure 5D, Table S6). Overall, approximately 31% of FERM species in the discovery cohort exhibited significant imbalance, a proportion that increased to 81% in the validation cohort (Figure 5E). These results indicate a consistent and widespread functional imbalance of FERM species across independent cohorts.

### Hypertension Is Associated with Increased Gut Microbial Metabolic Activity and Flux Intensification

To investigate the potential impact of gut microbiota dysbiosis on hypertension via microbial cross-feeding mechanisms, we further analyzed the metabolic fluxes of each metabolite by quantifying their production and consumption intensities within the cross-feeding context. After adjusting for potential confounders, we observed that the majority of metabolites in the hypertension group exhibited increased production and uptake fluxes, suggesting an overall enhancement of microbial metabolic activity in hypertensive individuals. (Figure S3A, Table S7) Furthermore, the adjusted average metabolic fluxes showed no significant correlation with *α*-diversity or microbial load but was positively correlated with microbial load. (Figure S3B.) By analyzing the fluxes of microbial substrate consumption and product synthesis, we investigated differences in gut microbial metabolic activity between patients with hypertension and healthy controls. The results revealed significant alterations in both uptake and production fluxes across multiple key metabolic pathways. In particular, pathways related to amino acid metabolism, carbohydrate metabolism, central carbon metabolism, polysaccharide metabolism, and inorganic nutrient metabolism exhibited enhanced activity under hypertensive conditions. (Figure S3C, Table S8) These findings suggest that hypertension is associated with a systematic reprogramming of gut microbial metabolic functions. Notably, enrichment analysis of metabolite production fluxes revealed increased activity of the butyrate metabolism pathway in patients with hypertension. Among the metabolites produced, butyrate showed a non-linear trend with increasing blood pressure, initially rising and then decreasing (Figure S3D).

## Discussion

This study systematically investigated alterations in gut microbiota composition, metabolic activity, and interspecies interactions under hypertensive conditions. By integrating microbial abundance with community-scale flux modeling, we identified multi-level disruptions in the gut microbial ecosystem, providing evidence for its role in hypertension pathogenesis.

Analysis of *α*-diversity showed significantly reduced microbial richness (Observe, Chao1, ACE) in hypertensive individuals, while evenness was unchanged. This suggests a loss of unique microbial taxa under hypertensive conditions. Declines in microbial richness are a hallmark of gut dysbiosis and have been widely reported in various metabolic and cardiovascular diseases [17]. From a taxonomic perspective, LDA identified multiple differentially abundant genera in the hypertensive group. This trend implies that hypertension may be associated with the loss of potentially protective or regulatory microbial taxa, which may in turn impair the functional capacity of the gut microbiota to contribute to host blood pressure homeostasis. *β* diversity analysis showed that the variance explained by the principal coordinates in the PCoA was low, and the *R*^2^ value from PERMANOVA was only 0.01, suggesting minimal overall structural differences in the gut microbiota between the hypertensive and control groups. These results highlight the need to investigate microbial interactions and metabolic fluxes to better understand hypertension-related dysbiosis.

Investigation of community-level medium metabolites identified 17 metabolites associated with hypertension, many of which have been previously reported to possess proinflammatory, anti-inflammatory, antioxidant properties, or function as cofactors for blood pressure-lowering agents. Among them, meso-2,6-diaminoheptanedioate, L-glutamate and L-cysteinylglycine have demonstrated the potential to directly or indirectly promote hypertension through mechanisms such as inflammation and accumulation of reactive oxygen species. In contrast, magnesium, cellobiose, Dextran 40, 5,6,7,8-tetrahydrofolate, amylopectin, melibiose, pyridoxine, 4-hydroxyphenylacetate, and carbon dioxide have been shown to exert antioxidant effects or promote the production of short-chain fatty acids or hypotensive factors. Pyridoxal 5-phosphate and L-serine have exhibited antioxidative or antihypertensive potential in some studies, though other evidence suggests they may also contribute to elevated blood pressure under certain conditions. Kestopentaose has been reported to enhance short-chain fatty acid synthesis, although a few studies found no significant effect. Octadecanoate (n-C18:0) has predominantly been associated with pro-inflammatory effects, which may be linked to compensatory microbial responses in the gut (Table S9). Most existing studies rely on plasma or serum-based metabolomic analyses [10], whereas this study employs a metabolic flux simulation of the gut microbiota to explore microbial metabolic capacity. As such, it offers a novel perspective on the pathophysiological mechanisms of hypertension from the standpoint of microbial functional metabolism.

Although cross-feeding networks showed no differences in node number or density, the hypertension group had shorter path lengths, suggesting more efficient metabolite transfer. Most metabolites exhibited positive effects on blood pressure (DoubleML), and this restructuring was unrelated to *α*-diversity. This change cannot be simply interpreted as a compensatory mechanism for reduced microbial diversity; species richness is not a key driver of metabolic network restructuring. Instead, this structural alteration is more likely driven by functional redundancy, regulation capacity of key metabolic pathways, or the ecological niche adaptability of certain dominant species, rather than solely relying on overall species count.

Hypertensive individuals’ gut microbiota show enhanced metabolic activity in pathways including methionine/cysteine, alanine/aspartate, the urea cycle, sulfur, and butyrate metabolism, which are central to microbial energy production and linked to host oxidative stress and signaling molecules such as *NO* and *H*_2_*S* [24, 8, 21]. In individuals with hypertension, the methionine and cysteine metabolic pathway is upregulated, which may affect the levels of homocysteine and *H*_2_*S*, thereby influencing vascular function [40]. Gut microbiota can convert methionine into homocysteine, and elevated homocysteine levels may promote oxidative stress by inhibiting *NO* synthesis [32]. In contrast, microbiota-derived *H*_2_*S* exerts vasodilatory effects [48]. Alanine and aspartate can participate in the urea cycle and one-carbon metabolism, providing precursors for *NO* synthesis [3]. In addition, the upregulation of microbial respiratory chain genes enhances the microbiota’s capacity to produce reactive oxygen species (ROS). Trace amounts of ROS or oxidative metabolites may translocate across the intestinal mucosa, inducing low-grade oxidative stress in the host [29, 33]. Notably, butyrate metabolism was enhanced in the hypertension group, characterized by an increased production rate of butyrate. As a key short-chain fatty acid, butyrate exhibits multiple physiological functions, including anti-inflammatory effects, vascular protection, and maintenance of intestinal barrier integrity [29, 28]. Butyrate flux is initially elevated, possibly as a compensatory response, but declines with progression to grade 2 hypertension, suggesting potential feedback inhibition or metabolic imbalance and warranting further investigation. Moreover, network module analysis revealed that the disruption of internal interactions within the FERM guild is associated with hypertension. The guild includes *Faecalibacterium, Enterobacter*, and *Roseburia* are anaerobic bacteria. Previous studies have demonstrated that Faecalibacterium produces SCFAs, such as butyrate, through the fermentation of dietary fiber [5], while *Enterobacter* is a Gram-negative bacterium capable of producing endotoxins such as lipopolysaccharides (LPS), which can trigger host immune responses and lead to inflammation [7]. Therefore, *Enterobacter* may contribute to hypertension by producing increased levels of endotoxins, which elicit host immune responses and promote inflammation. *Roseburia* produces butyrate and other SCFAs through the fermentation of complex polysaccharides [23], supporting intestinal barrier integrity and exerting anti-inflammatory effects. Although *Roseburia* has been implicated in hypertension-related studies [39]), our study did not observe significant differences in its abundance. However, its interaction patterns with other species were notably altered. Recent studies have shown Even without changes in species abundance, microbial network edge weights can be reorganized in disease, reflecting altered cooperation and competition that affect functional potential. [9]. Furthermore, using data from two cohorts, we found that the contribution of the FERM genus to key metabolites was significantly associated with blood pressure, and there was also a clear imbalance between the FERM genus and other species.

Based on these findings, we propose the following hypothesis: in hypertension, the gut microbial ecosystem exhibits metabolic dysregulation, characterized primarily by enhanced metabolic activity. This is reflected in the upregulation of pathways such as methionine and cysteine metabolism, the urea cycle, and alanine and aspartate metabolism, which may impair the microbial production of vasoregulatory molecules such as *NO* and *H*_2_*S*, thereby influencing blood pressure regulation. In parallel, the interaction network among gut microbes is substantially altered, particularly within the FERM microbial guild, whose disrupted associations are closely linked to 17 hypertension-related metabolites identified via DoubleML analysis. For example, L-glutamate may modulate blood pressure through neural regulatory mechanisms; 4-hydroxyphenylacetate may promote hypertension by activating pro-inflammatory signaling pathways; fermentation of cellobiose, dextran 40, and amylopectin generates SCFAs, which may influence blood pressure by modulating SCFA synthesis and signaling; meso-2,6-diaminoheptanedioate may activate NOD1 receptors in intestinal epithelial cells, triggering immune responses that impact vascular function. Additionally, 5,6,7,8-tetrahydrofolate, pyridoxine, and magnesium may participate in blood pressure regulation through their involvement in *NO* biosynthesis (Figure S4.).

This study proposes several mechanistic hypotheses regarding the relationship between gut microbiota metabolic functions and hypertension, providing potential insights for gut-targeted interventions or therapeutic strategies for hypertension. However, several limitations should be noted. First, the study is based on cross-sectional data, which limits the ability to draw definitive causal inferences. Second, the metabolic flux simulations depend on the accuracy of gut microbial abundance measurements and the completeness of metabolic models in the database. Additionally, dietary information from the samples was not collected; however, since all samples originated from Colombia, Western diet assumptions were uniformly applied as dietary constraints for metabolic flux prediction. Despite these limitations, this study lays a foundation for future mechanistic validation and intervention research on the gut microbiota–hypertension axis. Subsequent studies will integrate longitudinal cohort data, targeted metabolomics, and intervention trials to further validate these key findings and explore the feasibility of modulating the gut microbiome to improve blood pressure regulation.

## Conclusion

This study integrates gut microbial composition, metabolic flux, and microbial interaction networks to reveal structural and functional disturbances of the gut microbiota in hypertension. Hypertensive individuals showed reduced microbial *α*-diversity, especially in species richness, suggesting a loss of key functional taxa. Differential abundance and LDA analyses confirmed significant community shifts.

Metabolic flux analysis identified 17 metabolites associated with blood pressure regulation, involving inflammation, oxidative stress, and neuroactive pathways. Some metabolites (e.g., meso-2,6-diaminoheptanedioate, L-glutamate) were potentially pro-hypertensive, while others (e.g., magnesium, cellobiose, tetrahydrofolate) showed antihypertensive and antioxidant effects. Network module analysis revealed that the contribution of the FERM genus to key metabolites is significantly associated with blood pressure, alongside an imbalance between FERM genus and other microbes.

## Data availability

The species abundance tables used in this study originate from the dataset published by Jacobo de la Cuesta-Zuluaga et al., which is accessible at https://github.com/jsescobar/westernization. The data and code used in this study can be accessed in the GitHub repository at https://github.com/as147596/HT

## Author contributions statement

W.L. and Y.Z. writed original draft, data curation, validation, analysis, visualization. F.L., S.H. and S.L. writed review and editing, Methodology, Supervision. F.Y., S.S. and Z.W. managed this project, writed review and editing. F.Y. acquired funding, managed this project and writed original draft.

## Acknowledgments

We thank National Natural Science Foundation of China [62102065], the Fujian Provincial Health Commission [2022ZD01003], Fujian Medical University Research Foundation of Talented Scholars [XRCZX2022003] and Taizhou Science and Technology Plan Project [24ywa61] provided the financial support.

